# Cellular Interplay and Cytokine Hierarchy Cause Pathological Cardiac Hypertrophy in *RAF1*-Mutant Noonan Syndrome

**DOI:** 10.1101/122150

**Authors:** Jiani C. Yin, Mathew J. Platt, Xixi Tian, Xue Wu, Peter H. Backx, Jeremy A. Simpson, Toshiyuki Araki, Benjamin G. Neel

**Affiliations:** Department of Medical Biophysics, University of Toronto, Toronto, ON, M5G 1L7, Canada; Princess Margaret Cancer Centre, University Health Network, Toronto, ON, M5G 1L7, Canada; Laura and Isaac Perlmutter Cancer Center, New York University Langone Medical Center, New York, NY, 10016, USA; Department of Human Health and Nutritional Sciences, University of Guelph, Guelph, ON, N1G 2W1, Canada; Department of Biology, York University, Toronto, ON, M3J 1P3, Canada

**Author notes:** **Current address:** Geneseeq Technology Inc., Toronto, ON, M5G 1L7, Canada. Correspondence should be addressed to J.A.P., T.A., or B.G.N.

## Abstract

Noonan syndrome (NS) is caused by mutations in RAS/ERK pathway genes, and is characterized by craniofacial, growth, cognitive and cardiac defects. NS patients with kinase-activating *RAF1* alleles typically develop pathological left ventricular hypertrophy (LVH), which is reproduced in *Raf1^L613V/+^* knock-in mice. Here, using inducible *Raf1^L613V^* expression, we show that LVH results from the interplay of cardiac cell types. Cardiomyocyte *Raf1^L613V^* enhances Ca^2+^ sensitivity and cardiac contractility without causing hypertrophy. *Raf1^L613V^* expression in cardiomyocytes or activated fibroblasts exacerbates pressure overload-evoked fibrosis. Endothelial/endocardial (EC) *Raf1^L613V^* causes cardiac hypertrophy without affecting contractility. Co-culture and neutralizing antibody experiments reveal a cytokine (TNF/IL6) hierarchy in *Raf1^L613V^-expressing* ECs that drives cardiomyocyte hypertrophy *in vitro*. Furthermore, post-natal TNF inhibition normalizes the increased wall thickness and cardiomyocyte hypertrophy *in vivo*. We conclude that NS cardiomyopathy involves cardiomyocytes, ECs, and fibroblasts, TNF/IL6 signaling components represent potential therapeutic targets, and abnormal EC signaling might contribute to other forms of LVH.

## Introduction

Pathological left ventricular hypertrophy (LVH) is a common inherited disorder (∼1 in 500 live births), and represents the leading cause of sudden death in young people^1–3^. Features of pathological hypertrophy include increased cardiomyocyte (CM) size, thickening of the ventricular wall and septum, perivascular and interstitial fibrosis, and cardiac dysfunction that can eventuate in heart failure^2,3^. LVH has many potential etiologies, including hypertension, cardiac valve disease or genetic defects^4–6^. Most inherited forms of LVH are caused by mutations in genes encoding sarcomeric proteins^1,3,5,7^; hence, most studies of these disorders have focused on the CM-intrinsic effects of these genes. However, ∼25% of cases are caused by mutations in genes that encode signal transduction components. Genetic analyses and transgenic animal models have confirmed that aberrant signaling can drive pathological cardiac hypertrophy, often in association with other systemic defects^8–11^.

Abnormal regulation of the RAS/RAF/MEK/ERK (hereafter, RAS/ERK) pathway underlies a group of related developmental syndromes termed “RASopathies”, which are characterized by a spectrum of phenotypes, including craniofacial dysmorphia, delayed growth, cognitive problems and cardiac abnormalities^12–14^. The most common RASopathy, Noonan Syndrome (NS), can be caused by germ-line gain-of-function *PTPN11, KRAS, NRAS, RRAS, SHOC2, SOS1/2, RAF1, RIT1* or *PPP1CB* alleles^12^,^15–20^. Pathological LVH (NS-cardiomyopathy) is seen in ∼20% of NS cases overall. Kinase activating mutations in *RAF1*, which encodes a serine/threonine kinase for MEK, account for a small fraction (∼5%) of NS, but nearly all (>95%) such patients develop NS-cardiomyopathy^21^. Previously, we found that *Raf1^L613V/+^* knock-in mice recapitulate the human disorder, with growth defects, facial dysmorphia, and most importantly, eccentric LVH^22^. Specifically, in *Raf1^L613V/+^* knock-in mice, heart mass, ventricular wall thickness, ventricular chamber dimensions and cardiac contractility are increased, and cardiac fibrosis following pressure overload is exacerbated.

Capitalizing on our inducible *Raf1^L613V^* allele (*Raf1^L613Vfl^*)*^22^*, we deconstructed the above cardiac phenotypes into their contributing cell types. Surprisingly, although expression of mutant *Raf1* in CMs caused altered contractility as a consequence of increased calcium sensitivity, it did not result in pathological LVH. Mutant expression in activated cardiac fibroblasts or CMs resulted in an increased fibrotic response to pressure overload. Intriguingly, mutant expression in ECs, by means of a TNF/IL6 cytokine hierarchy, led to increased CM and chamber size without affecting contractility. Treatment of *Raf1* mutant mice with anti-TNF antibodies reversed their increased CM size and wall thickness. Our results reveal the cellular and molecular complexity underlying NS cardiomyopathy and suggest that anti-TNF antibodies could be a therapeutic option for severe pathological LVH in NS patients.

## Results

### CM-specific *Raf1^L613V^* expression alters contractility

We crossed *Raf1^L613Vfl/+^* knock-in mice with lineage-specific Cre recombinase lines, which catalyzed “STOP” cassette deletion selectively in the expected cell types (Fig. 1a and Supplementary Fig. 1a). Mutant expression in CMs was achieved using mice expressing Cre under the control of the *Mlc2v* promoter, which is activated efficiently and exclusively in ventricular CMs as early as E8.75^23^. As anticipated, the “STOP” cassette in the *Raf1^L613V^* allele had been excised efficiently in the hearts of 10 week-old mice (compare to global *Raf1^L613V/+^* mice; Supplementary Fig. 1b), as well as in isolated CMs at post-natal day 4 (Supplementary Fig. 1c). Surprisingly, and in stark contrast to global *Raf1^L613V/+^* mice, heart weight to body weight (HW/BW) ratio and CM cross-sectional area in Mlc2v-L613V mice were comparable to littermate controls (Fig. 1b, c). Similarly, HW/BW ratio was not increased (and in fact, was slightly decreased) when *Raf1^L613V^* expression was induced in atrial and ventricular myocardium under the control of the cardiac troponin T promoter^24^ (Supplementary Fig. 1d). Echocardiography, performed at 16 weeks of age, revealed reduced left ventricular internal end-systolic (LVIDs) and -diastolic (LVIDd) dimensions (Fig. 1d, e) and a small increase in left ventricular posterior wall thickness (LVPWd; Fig. 1f) in Mlc2v-L613V hearts.

**Figure 1.**
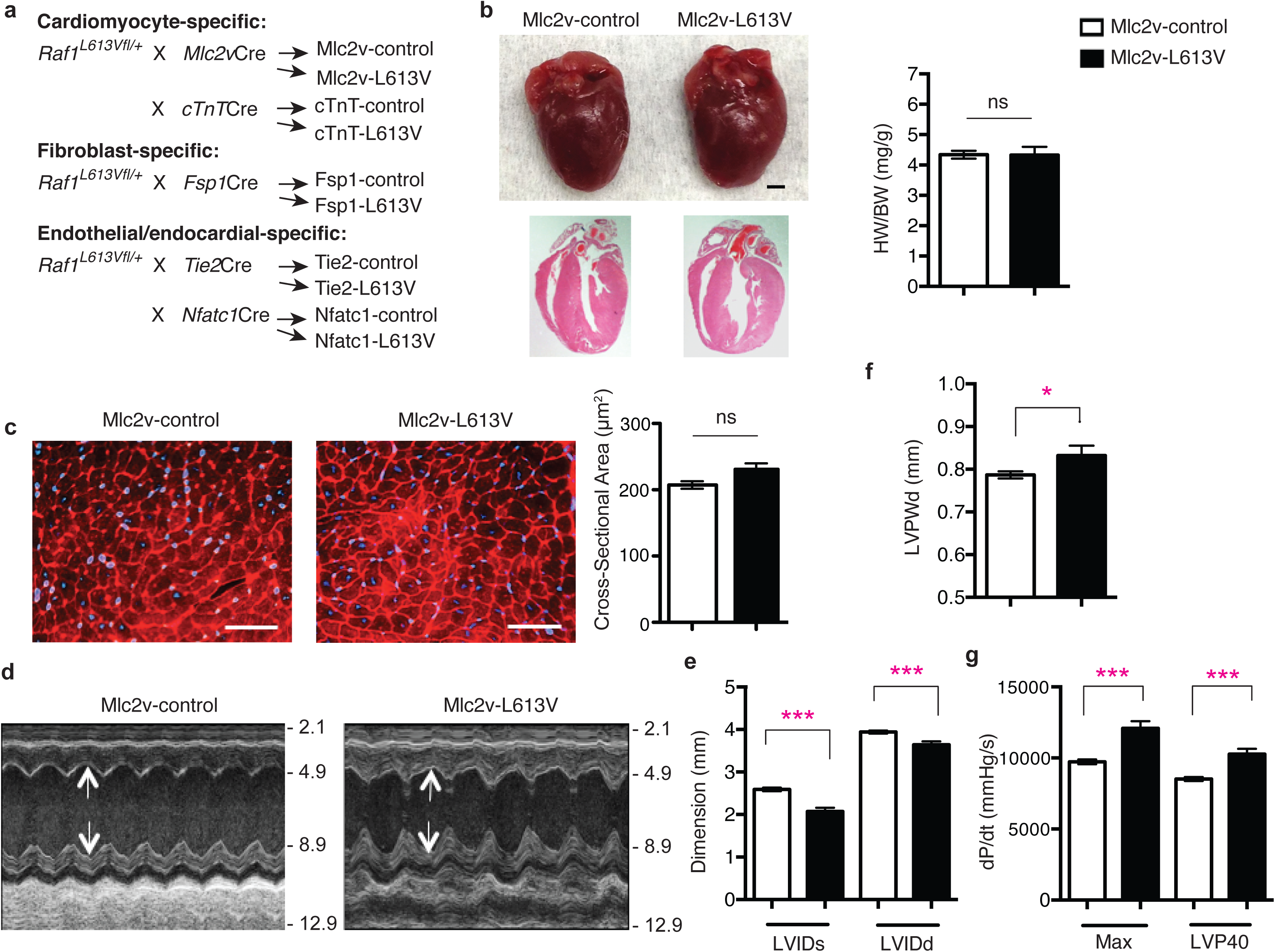
CM-specific *Raf1^L613V^* expression causes enhanced contractility. **a,** Schemes for generating mice with tissue-specific *Raf1^L613V^* expression and controls (controls are a combination of wild type, *Raf1^L613Vfl/+^*, and the respective Cre mice, unless indicated otherwise; no significant differences were seen between control groups across all experiments). **b**, Representative gross morphology and H&E-stained sections of Mlc2v-L613V and littermate control hearts at week 10 (scale bar: 1 mm). Heart weight to body weight (HW/BW) was measured at 4 months of age (mean ± S.E.M.; *n* = 38 (Mlc2v-control) or 13 (Mlc2v-L613V)). **c**, Representative wheat germ agglutinin (WGA)-stained cross-sections of hearts from 10-week old Mlc2v-L613V and control mice (original magnification, ×400; scale bars: 50 μm). Cross-sectional area (right) was quantified using ImageJ (mean ± S.E.M.; *n* = 5 samples per group, with 200 CMs counted per sample; *P* = 0.18, two-tailed Student’s *t* test). **d**, Representative echocardiograms of hearts from 4-month-old mice. Arrows indicate LV diastolic dimensions. **e**, Left ventricular end-systolic and end-diastolic dimensions (LVIDs and LVIDd, respectively) and **f**, Left ventricular posterior wall thickness (LVPWd), measured by echocardiography at 4 months (mean ± S.E.M.; *n* = 43 (Mlc2v-control) or 17 (Mlc2v-L613V); \**P* < 0.05, \*\*\**P* < 0.001, two-tailed Student’s *t* test). **g**, Cardiac contractility of 4-month old mice, measured by invasive hemodynamics (mean ± S.E.M.; *n* = 39 (Mlc2v-control) or 16 (Mlc2v-L613V). *\*\*\*P* < 0.0001, two-tailed student’s *t* test).

Although Mlc2v-L613V hearts showed minimal hypertrophy, they were markedly hyper-contractile. As in global *Raf1^L613V/+^* mice, ejection fraction (EF) and fractional shortening (FS) were increased in Mlc2v-L613V mice, compared with controls (Supplementary Table 1). Invasive hemodynamics revealed increased *dP/dt_max_* and *dP/dt@*LVP40 (Fig. 1g); the latter is independent of the slightly reduced afterload (systolic pressure) observed in these mice (Supplementary Table 2). Thus, CM-restricted RAF1 mutant expression induces a hyper-contractile state in the absence of CM hypertrophy.

To ask whether the effects of the RAF1 mutant on cardiac contractility originated from alterations in Ca^2+^ homeostasis, we loaded CMs isolated from Mlc2v-control or -L613V mice with Fura-2 and measured Ca^2+^ transients (i.e., R_340/380_) in response to field stimulation at 0.5 Hz. Representative Ca^2+^ transients were similar in freshly isolated single Mlc2v-control and -L613V CMs (Supplementary Fig. 2a). Neither basal Ca^2+^ levels, nor the Ca^2+^ transient peaks, differed between the groups (Supplementary Fig. 2b: basal R_340/380_ was 1.00 ± 0.02 for Mlc2v-control vs. 1.03 ± 0.02 for Mlc2v-L613V, *P* = 0.24, Student’s *t* test; peak R_340/380_ was 1.82 ± 0.05 for Mlc2v-control vs. 1.81 ± 0.05 for Mlc2v-L613V, *P* = 0.90, Student’s *t* test). Likewise, no differences were observed in the kinetics of the Ca^2+^ transients (Supplementary Fig 2c: time to the R_340/380_ peak 47.72 ± 2.57 ms in Mlc2v-control vs. 46.12 ± 2.36 ms in Mlc2v-L613V, *P* = 0.65, Student’s *t* test; time to 50% decay of the R_340/380_ was 218.17 ± 8.07 ms for Mlc2v-control vs. 210.62 ± 9.36 ms for Mlc2v-L613V, *P*=0.55, Student’s *t* test). Quantitative reverse-transcription PCR (qRT-PCR) and immunoblots of lysates from Mlc2v-L613V hearts showed similar levels of SERCA mRNA and protein, respectively, as in littermate control hearts (Supplementary Fig. 2d, e and 9).

Enhanced cardiac contractility in Mlc2v-L613V mice in the absence of elevated intracellular Ca^2+^ levels might reflect increased Ca^2+^-sensitivity of the contractile apparatus. We assessed Ca^2+^ sensitivity by determining the relationship between force and Ca^2+^ during twitches induced by field stimulation of isolated CMs attached to stiff glass rods (Supplementary Fig. 3)^25,26^. Plots of force as a function of Ca^2+^ (phase plots^25^) during the very late phase of the force relaxation curve became independent of peak force levels when CMs were held at a resting sarcomere length of 1.8 μm (Supplementary Fig. 4). Hence, the force-Ca^2+^ relationship achieves a steady state during the terminal phase of relaxation, which enables assessment of the Ca^2+^ sensitivity of CM contractile proteins^25^. Typical simultaneous force-Ca^2+^ recordings for Mlc2v-control and -L613V CMs revealed comparable decay kinetics of the Ca^2+^ transient (solid lines) in both groups, but the force in the late phase of relaxation (dashed lines) was elevated markedly in Mlc2v-L613V CMs, suggesting higher Ca^2+^ sensitivity (Fig. 2a, b). Superimposing the force-Ca^2+^ phase plots for several Mlc2v-control and -L613V CMs confirmed that the force in mutant CMs was generally above the force generated by control CMs during the late phase of the twitch relaxation (Fig. 2c, d). The difference between control and mutant CMs is more obvious in Fig. 2e, which shows the force-Ca^2+^ relationship (insets in Fig 2c, d) on an expanded scale. The force averages at five Ca^2+^ levels for the same Mlc2v-control and -L613V CMs show that mutant CMs develop more force as a function of Ca^2+^; i.e., that the calcium sensitivity of their contractile proteins is increased (Fig. 2f).

**Figure 2.**
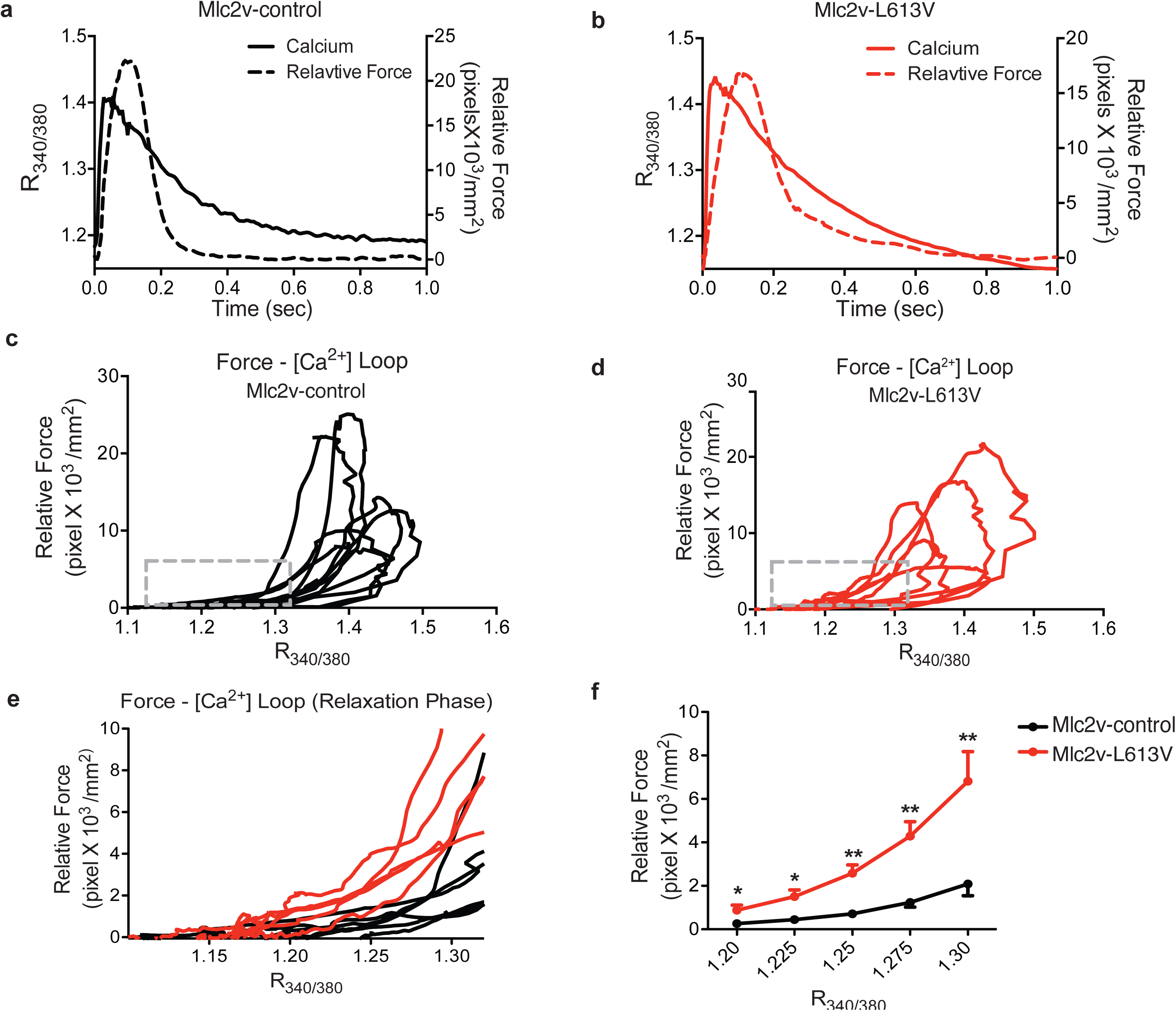
Increased Ca^2+^ sensitivity in Mlc2v-L613V hearts. Force-Ca^2+^ relationships measured in single isolated CMs during twitches in response to field stimulation. **a** and **b**, Typical Ca^2+^ transients and force measurements for Mlc2v-control (**a**) and -L613V (**b**) myocytes attached to glass rods. Note that during the late phase of relaxation, the force level is much higher in the Mlc2v-L613V CMs compared with controls, despite their similar Ca^2+^ transients. **c** and **d**, Superimposed force-Ca^2+^ relationships in response to field stimulation (i.e., phase plots^25^) for several Mlc2v-control (**c**) and -L613V (**d**) CMs. **e**, Magnified and superimposed force/Ca^2+^ recordings for Mlc2v-control (black) and -L613V (red) CMs during late phase relaxation, when the relationship between force and Ca^2+^ reaches a steady state (Supplementary Fig. 4). **f**, Averaged force levels at 5 selected Ca^2+^ levels, showing upward shift of Mlc2v-L613V, compared with control, CMs, indicative of greater force generation for the same Ca^2+^ level (* *P* < 0.05, ** *P* < 0.01, two-tailed Student’s *t* test).

### RAF1 mutant CMs and activated fibroblasts increase fibrosis

The above results imply that (an)other cardiac cell type(s), via paracrine signaling, promotes hypertrophy in RAF1-mutant NS. At baseline, no difference was detected in cardiac morphology or contractile function, save for a slight reduction in LVPWd, in Fsp1-L613V mice, compared with their respective controls (Fig. 3a-d, Supplementary Table 1, 2). However, expression of the mutant *Raf1* allele was barely detectable in Fsp1-L613V hearts (Supplementary Fig. 5a), consistent with the greater activity of the Fsp1-Cre line in activated fibroblasts^27,28^, but making it difficult to reach strong conclusions about the effects of mutant expression in fibroblasts under basal conditions. To gain some insight into this issue, we asked whether RAF1 mutant CFs can promote CM hypertrophy *in vitro*. To this end, we co-cultured CD90^+^ cells from *Raf1^L613V/+^* global knock-in hearts^22^ with wild type CMs. Notably, myocyte size was comparable to those co-cultured with control CD90^+^ cells (Supplementary Fig. 5b); under the same conditions, mutant ECs promote CM hypertrophy *in vitro* (see below).

**Figure 3.**
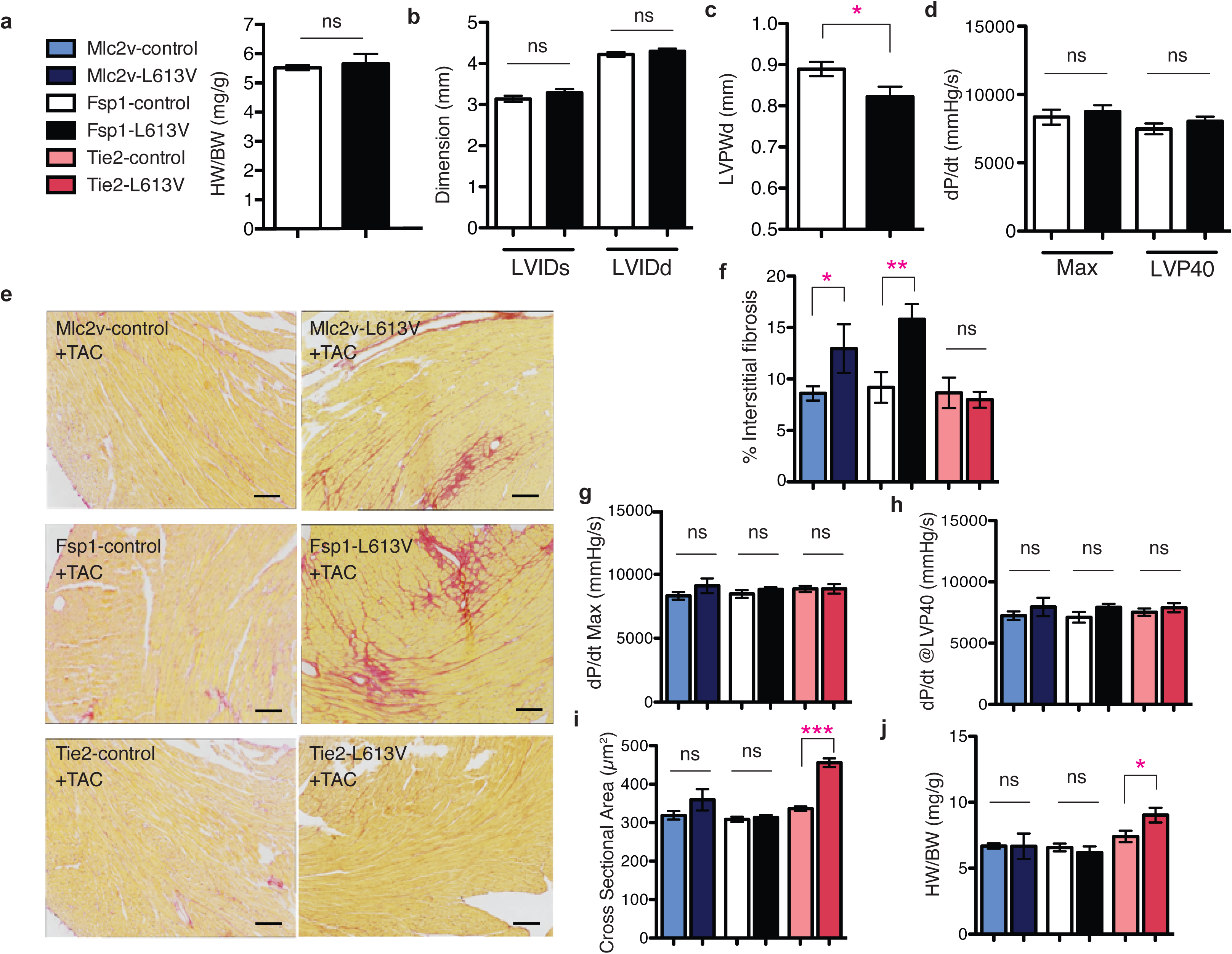
CM- or *Fsp1-driven Raf1^L613V^* expression enhances fibrosis after TAC. **a-d,** Baseline measurements. **a**, Baseline HW/BW ratio of Fsp1-L613V and control mice at 10 weeks of age. **b**, Baseline LVIDs and LVIDd and **c**, LVPWd, as measured by echocardiography at 4 months (mean ± S.E.M.; *n* = 18 (Fsp1-control); 7 (Fsp1-L613V); \**P* < 0.05, two-tailed Student’s *t* test). **d**, Baseline cardiac contractility of 4-month old mice, measured by invasive hemodynamics (mean ± S.E.M.; *n* = 9 (Fsp1-control) or 5 (Fsp1-L613V)). **e-j**, measurements at 2 weeks after TAC. **e**, Interstitial fibrosis in Mlc2v-L613V and Fsp1-L613V hearts, shown by Picro Sirius Red (PSR) staining; original magnification, ×100; scale bars: 100 μm) at 2 weeks after TAC. **f**, percent pixels staining positive for interstitial fibrosis with PSR were quantified using ImageJ (mean ± S.E.M.; *n* = 9 (Mlc2v-control), 4 (Mlc2v-L613V), 8 (Fsp1-control), 7 (Fsp1-L613V), 8 (Tie2-control) or 5 (Tie2-L613V); \**P* < 0.05, \*\**P* < 0.01, two-tailed Student’s *t* test). Note that there is no difference between Tie2-L613V and its respective controls. **g** and **h**, Cardiac contractility, measured by invasive hemodynamics 2 weeks post-TAC (mean ± S.E.M.; *n* = 9 (Mlc2v-control), 4 (Mlc2v-L613V), 9 (Fsp1-control), 8 (Fsp1-L613V), 36 (Tie2-control) or 11 (Tie2-L613V)). Note absence of contractility difference between Mlc2v-L613V and controls following 2 weeks of TAC. **i**, Cross-sectional area, quantified by using ImageJ (mean ± S.E.M.; *n* = 9 (Mlc2v-control), 4 (Mlc2v-LV), 9 (Fsp1-control), 8 (Fsp1-LV), 11 (Tie2-control) or 5 (Tie2-L613V), with 200 CMs counted per sample; \*\*\**P* < 0.0001, two-tailed Student’s *t* test). **j**, HW/BW ratio 2 weeks post-TAC (mean ± S.E.M.; *n* = 6 (Mlc2v-control), 2 (Mlc2v-LV), 9 (Fsp1-control), 3 (Fsp1-LV), 17 (Tie2-control) or 4 (Tie2-L613V); \**P* < 0.05, one-tailed Student’s *t* test).

FSP1 expression is induced in activated cardiac (and other) fibroblasts upon stress (Supplementary Fig. 5a)^27,28^. Indeed, compared with controls, Fsp1-L613V mice developed more severe interstitial fibrosis within two weeks of biomechanical stress imposed by transverse aortic constriction (TAC; Fig. 3e, f). Interestingly, the extent of the fibrotic response did not correlate with a more severe impairment in cardiac function, as EF, FS, *dP/dt_max_* and *dP/dt@*LVP40 were comparable in Fsp1-L613V and littermate control mice following TAC (Fig. 3g, h and Supplementary Table 1, 2). FSP1 also is reported to mark hematopoietic and endothelial cells, and endothelial-to-mesenchymal transition (EndoMT) can contribute to cardiac fibrosis^29,30^. However, Tie2-L613V mice, which direct *Raf1^L613V^* expression to endocardial/endothelial cells (ECs), did not develop more cardiac fibrosis than controls after TAC, arguing against a pro-fibrotic role for mutant expression in these cells (Fig. 3e, f). We cannot, however, exclude the less parsimonious possibility that dual action in ECs and activated fibroblasts mediates fibrosis in Fsp1-L613V mice.

Enhanced fibrosis also was observed in Mlc2v-L613V hearts in response to TAC (Fig. 3e, f), along with loss of their hyper-contractility (Fig. 3g, h and Supplementary Table 1, 2). Conceivably, prolonged pressure overload would lead to further deterioration of cardiac function, and ultimately, functional decompensation, in Mlc2v-L613V mice, as seen in global *Raf1^L613V/+^* mice at 8 weeks after TAC^22^. Notably, TAC did not alter the hypertrophic response in Mlc2v-L613V or Fsp1-L613V hearts compared to their respective controls (Fig. 3i, j).

### EC-specific *Raf1^L613V^* expression causes cardiac hypertrophy

By contrast, selective expression of *Raf1^L613V^* in ECs (achieved by crossing *Raf1^L613Vfl/+^* mice to *Tie2Cre* or *Nfatc1*Cre mice; Supplementary Fig. 1c) was associated with marked cardiac hypertrophy both basally (Fig. 4a, b) and following TAC (Fig. 3i, j), as indicated by significant increases in HW/BW ratio and CM cross-sectional area. Cardiac hypertrophy was detectable as early as post-natal day 4 (Supplementary Fig. 6a, b), but there was no difference in the number of BrdU^+^ (proliferation) or TUNEL^+^ (apoptosis) CMs in embryonic Tie2-L613V or Nfatc1-L613V hearts (Supplementary Fig. 6c, d). Valvuloseptal development and function were normal, as assessed by histology (Supplementary Fig. 6e) and by the absence of a significant pressure gradient across the aortic valve (Supplementary Table 2). Systemic arterial pressure was lower in Tie2-L613V or Nfatc1-L613V mice than in controls (Supplementary Table 2), excluding hypertension as the cause of cardiac hypertrophy. Echocardiograms revealed markedly increased LVIDs and LVIDd (Fig. 4c, d), along with increased LVPWd (Fig. 4e). Remarkably, contractility remained within normal limits, as assessed by echocardiography and invasive hemodynamic analysis (Fig. 4f and Supplementary Table 1, 2). Even when subjected to TAC, cardiac contractility in Tie2-L613V mice remained comparable to littermate controls (Fig. 3g, h and Supplementary Table 1, 2). Hence, hypertrophy, hyper-contractility and fibrosis in global *Raf1^L613V/+^* mice are separable, and must reflect distinct cellular and molecular mechanisms.

**Figure 4.**
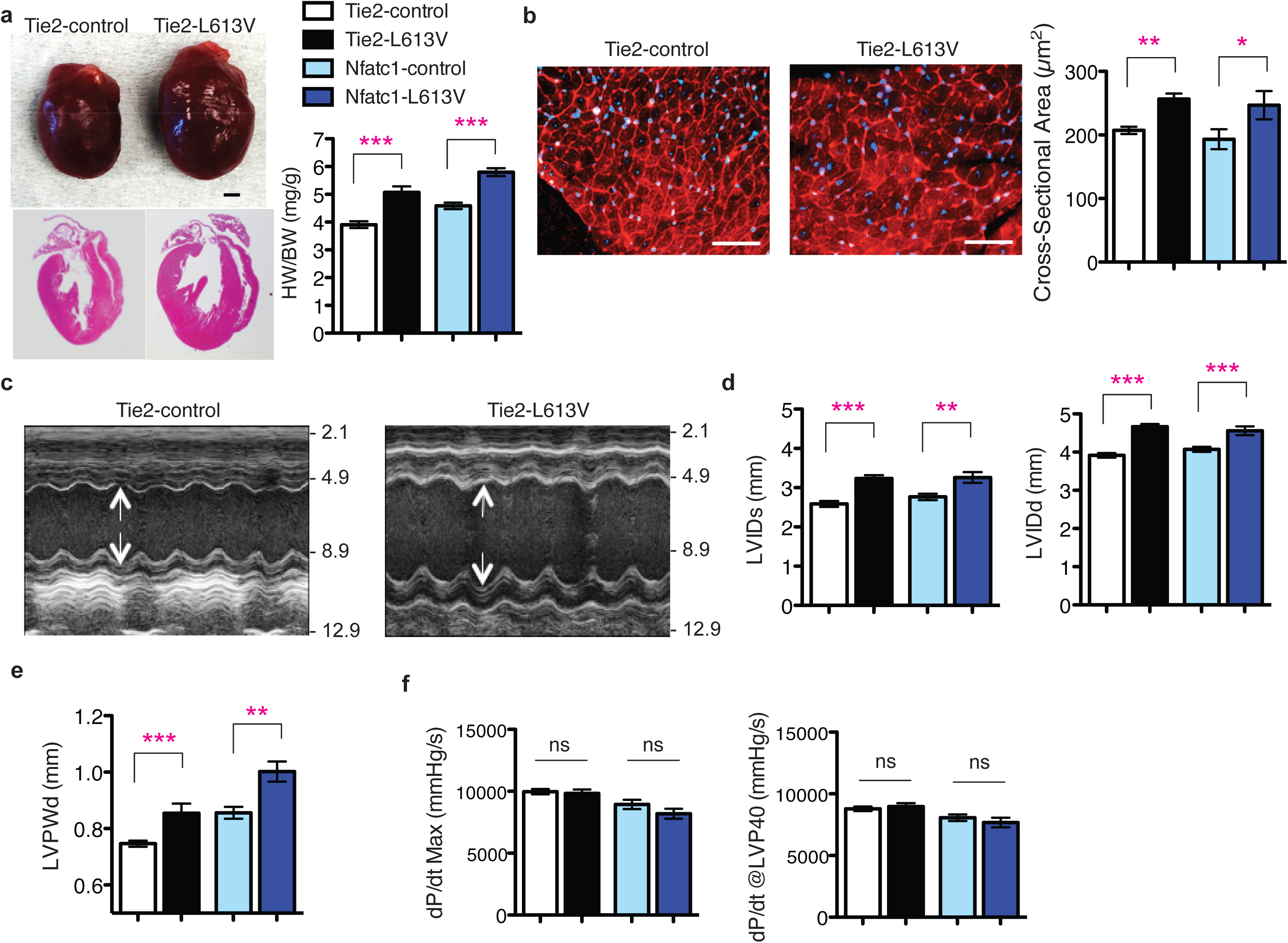
EC-specific *Raf1^L613V^* expression causes cardiac hypertrophy. **a,** Representative gross morphology and H&E-stained sections of Tie2-L613V and control hearts at 10 weeks (scale bar: 1 mm), and HW/BW of Tie2-L613V and Nfatc1-L613V mice, measured at 4 (mean ± S.E.M.; *n* = 27 (Tie2-control), 13 (Tie2-L613V), 15 (Nfatc1-control) or 8 (Nfatc1-L613V) months; \*\*\**P* < 0.0001, two-tailed Student’s *t* test). **b**, Representative WGA-stained cross-sections of 10-week old Tie2-L613V and control hearts (original magnification, ×400; scale bars: 50 μm). Cross-sectional area (right), quantified using ImageJ (mean ± S.E.M.; *n* = 5 samples per group, with 200 CMs counted per sample; \*\**P* < 0.005, \**P* < 0.05, two-tailed Student’s *t* test). **c**, Representative echocardiograms of hearts from 4 month-old mice. Arrows indicate LV diastolic dimensions. **d**, LVIDs and LVIDd and **e**, LVPWd, measured by echocardiography at 4 months (mean ± S.E.M.; *n* = 24 (Tie2-control), 12 (Tie2-L613V), 15 (Nfatc1-control) or 9 (Nfatc1-L613V); \*\**P* < 0.005, \*\*\**P* < 0.001, two-tailed Student’s *t* test). **f**, Cardiac contractility of 4 month-old mice, measured by invasive hemodynamics (mean ± S.E.M.; *n* = 27 (Tie2-control), 13 (Tie2-L613V), 15 (Nfatc1-control) or 8 (Nfatc1-L613V)).

### RAF1 mutant ECs induce CM hypertrophy via TNF/IL6 signaling

Endothelial cells in the heart can influence cardiac development and function via paracrine signals^31^. To explore the non-cell autonomous effects of endocardium/endothelium on CM hypertrophy, we purified RAF1 mutant or control cardiac endothelial/endocardial cells with CD31 magnetic beads, and co-cultured them with wild type neonatal CMs in serum-free media conditions (Fig. 5a). Consistent with our *in vivo* findings, CM surface area was increased after three days of co-culture with Tie2-L613V or Nfatc1-L613V ECs, compared with those co-cultured with controls (Fig. 5b and Supplementary Fig. 7a). RAF1 mutant CMs exhibited similar increases in cell size to control CMs after co-culture with RAF1 mutant ECs, confirming that signals emanating from ECs are the primary cause of CM hypertrophy (Supplementary Fig. 7b). A similar size increase occurred in Transwell assays (Fig. 5a, c), suggesting that (a) diffusible, paracrine factor(s) derived from RAF1 mutant ECs promotes CM hypertrophy.

**Figure 5.**
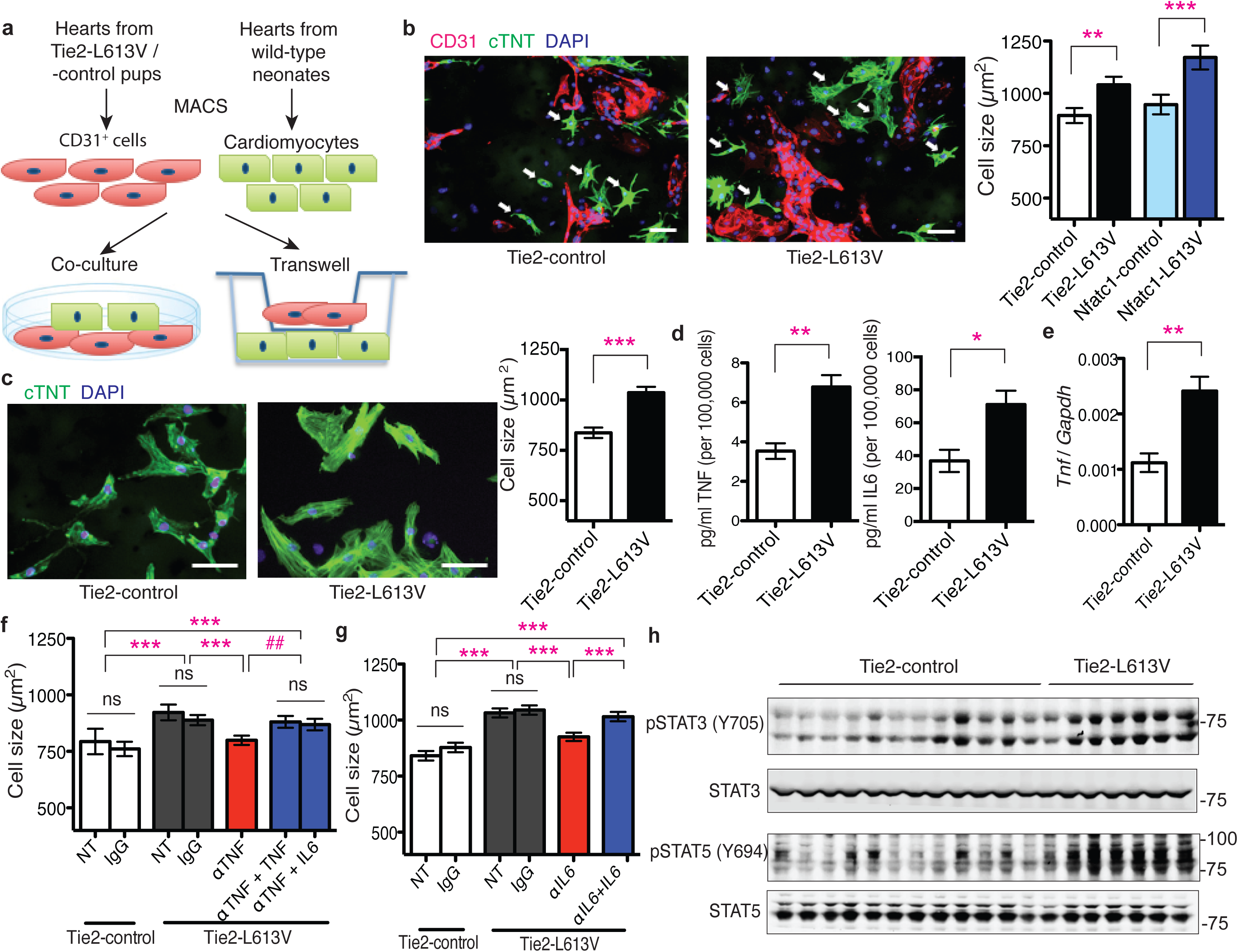
Cardiac ECs induce cardiomyocyte hypertrophy *in vitro*. **a,** Schematic of direct and Transwell co-culture experiments. **b**, Neonatal WT CMs were co-cultured for 3 days with CD31^+^ cells from post-natal day 4-7 Tie2-L613V or control hearts. Left: Representative immunofluorescence staining for CD31 (magenta), cardiac troponin T (green) and DAPI (blue) in co-cultures (original magnification, ×200, scale bar: 50 μm). Arrows indicate individual CMs that were among those assessed by ImageJ. Right: Quantification of one of three independent experiments with similar results. **c**, Left: Transwell assays. CD31^+^ cells were seeded in the upper chamber and sizes of neonatal WT CMs (green) in the bottom chamber were assessed using ImageJ (original magnification, ×400, scale bars: 50 μm). Right panel: Quantification of one of three independent experiments with similar results (mean ± S.E.M.; 300 CMs counted per group; \*\**P* < 0.005, *** *P* < 0.001, two-tailed Mann-Whitney test). **d**, Conditioned media were collected from CD31^+^ cultures two days after a media change, and TNF and IL6 levels were measured by Luminex assay (mean ± S.E.M.; *n* = 4 each group; \*\**P* < 0.005, \**P* < 0.05, two-tailed Student’s *t* test). **e**, *Tnf* mRNA levels in cultured ECs, assessed by qRT-PCR (mean ± S.E.M.; *n* = 4 each group; \*\**P* < 0.005, two-tailed Student’s *t* test). **f** and **g**, CMs and CD31^+^ cells were co-cultured overnight, and then subjected to either **f**, anti-TNF antibody (MP6-XT22, 1 ng/ml), or **g**, anti-IL6 neutralizing antibody (MP5-20F3, 10 ng/ml) or isotype control (IgG1) in the presence or absence of excess recombinant TNF (1 ng/ml) or IL6 (25 ng/ml), as indicated. Quantification is shown for one of three independent experiments with similar results (mean ± S.E.M.; 300 CMs counted per group; \*\**P* < 0.005, \*\*\**P* < 0.0001, Dunn’s post-hoc test when ANOVA (Kruskal-Wallis test) was significant; *^##^P* < 0.005, two-tailed Mann-Whitney test). Note that either exogenous TNF or IL6 reverses the inhibitory effect of anti-TNF antibody. **h**, Heart lysates from Tie2-L613V or control mice were analyzed by immunoblotting with the indicated antibodies. STAT3 and STAT5 levels serve as loading controls.

To search for this paracrine factor(s), we screened conditioned media from Tie2-L613V or control ECs for multiple cytokines, including known inducers of cardiac hypertrophy (Supplementary Fig. 7c). We observed 1.5-2 fold increases in the levels of Tumor Necrosis Factor alpha (TNF) and interleukin-6 (IL6) in conditioned media from mutant ECs (Fig. 5d). The *Tnf* promoter contains Activator protein-1 (AP-1) and ETS-1 (transcription factors downstream of ERK) binding sites^32,33^, and RNAseq analysis revealed increased transcripts of *Tnf* (among other genes) in mutant ECs (Supplementary Fig. 7d). We confirmed this finding by using qRT-PCR (Fig. 5e). By contrast, the increased IL6 production reflected post-transcriptional regulation (Supplementary Fig. 7e). The isolated CD31^+^ population contained almost no CD45^+^ cells (Supplementary Fig. 7f), and heart sections from Tie2-L613V mice exhibited comparable CD45 staining to their littermate controls (Supplementary Fig. 7g), arguing against contaminating hematopoietic cells as the source of the increased TNF and IL6 in EC-conditioned media.

Addition of recombinant mouse TNF or IL6 (at levels similar to those found in RAF1 mutant EC-conditioned media) induced dose-dependent increases in wild type CM size (Supplementary Fig. 8a, b). Furthermore, neutralizing anti-TNF or anti-IL6 monoclonal antibodies (but not cognate isotype-matched controls) blocked the pro-hypertrophic effects of RAF1 mutant ECs on co-cultured CMs. As expected, the effects of each neutralizing antibody were reversed by adding an excess of the cognate exogenous cytokine (Fig. 5f, g). These data argue that TNF/IL6 produced by mutant cardiac ECs play a critical role in the development of cardiomyopathy in RAF1-mutant NS.

In many inflammatory disorders, most notably, rheumatoid arthritis, TNF stands atop a cytokine hierarchy that can include IL6^34,35^. An analogous hierarchy appears to be present in RAF1-mutant ECs: TNF increased, whereas anti-TNF antibody treatment decreased the levels of IL6 in cardiac EC-conditioned media (Supplementary Fig. 8c). Furthermore, the hypertrophy-reducing effects of the neutralizing anti-TNF antibody were reversed by excess IL6, as well as by TNF, placing IL6 production “downstream” of TNF stimulation (Fig. 5f). IL6 signals via the IL6 receptor/gp130 complex, which in turn, activates the JAK/STAT, MEK/ERK and PI3K/AKT pathways. Consistent with IL6 acting as the proximate mediator of HCM *in vivo*, immunoblots of total heart extracts from Tie2-L613V mice, which express mutant RAF1 only in ECs but predominantly contain CM-derived proteins, revealed increased activation of STAT3 (phospho (p)-Tyr705) and STAT5 (p-Tyr694), MEK (p-Ser217/221), ERK (p-Tyr204Thr202) and AKT (p-Ser473 and p-Thr308; Fig. 5h, Supplementary Fig. 8d-I, 10 and 11), compared with control hearts.

### CM hypertrophy is normalized by TNF inhibition *in vivo*

Our co-culture experiments implicated EC-derived TNF in the pathogenesis of CM hypertrophy. We therefore asked if TNF inhibition could normalize EC-induced CM hypertrophy *in vivo*. Tie2-L613V or littermate control mice (4 week-old) were injected i.p. with anti-TNF neutralizing antibody or isotype control (5 mg/kg body weight) twice weekly for 6 weeks (Fig. 6a). Ventricular chamber dimensions and HW/BW ratio remained elevated in Tie2-L613V mice after 6 weeks of anti-TNF antibody treatment (Supplementary Fig. 8j-l). However, LVPWd was significantly reduced in hearts from anti-TNF antibody-treated Tie2-L613V mice, compared with isotype control-treated mice (Fig. 6b). We also observed a corresponding decrease in CM cross-sectional area in Tie2-L613V mice subjected to TNF inhibition (Fig. 6c). Notably, hearts from wild type littermates were not affected by anti-TNF antibody treatment, indicating that antibody effects are RAF1 mutant-specific. Furthermore, cardiac function was preserved in Tie2-L613V mice subjected to anti-TNF antibody therapy (Fig. 6d, e). Interestingly, the decreased blood pressure in Tie2-L613V mice also was normalized by anti-TNF treatment (LVP was 97.3 ± 2.4 mmHg in Tie2-L613V mice with isotype control treatment vs. 107.9 ± 0.9 mmHg in those with TNF inhibition).

**Figure 6.**
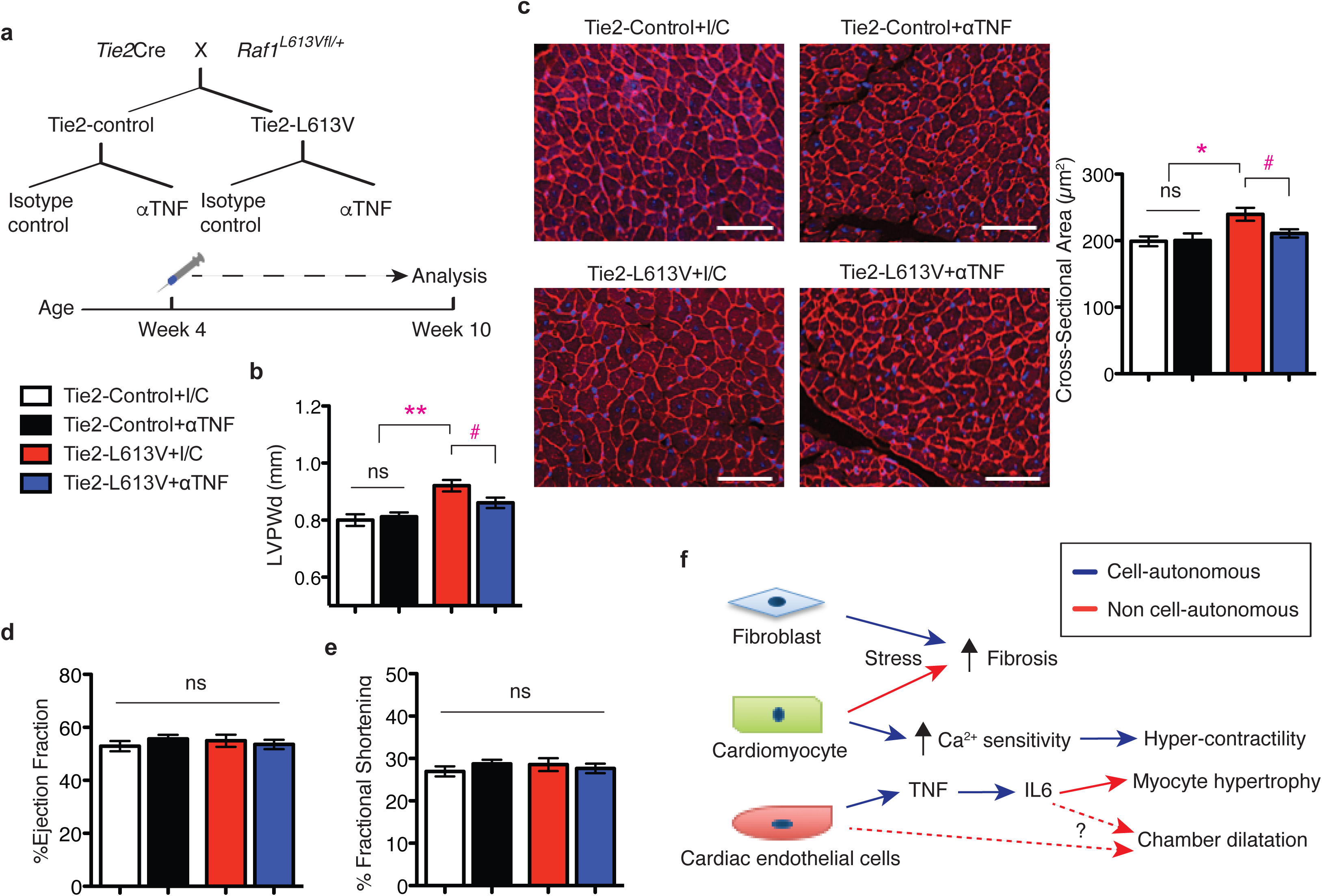
Anti-TNF antibody treatment normalizes CM hypertrophy *in vivo*. **a,** Schematic of the treatment regimen. **b**, TNF inhibition normalizes LVPWd after treatment for 6 weeks, as assessed by echocardiography (mean ± S.E.M.; *n* = 10 (Tie2-control + isotype control), 11 (Tie2-control + anti-TNF Ab), 7 (Tie2-L613V + isotype control) or 9 (Tie2-L613V + anti-TNF Ab); \*\**P* < 0.005, Bonferroni’s post-test when ANOVA was significant; # *P* < 0.05 two-tailed Student’s *t* test). **c**, Representative WGA-stained cross-sections of hearts from treated animals (original magnification, ×200; scale bars: 50 μm). Cross-sectional area (right), quantified using ImageJ (mean ± S.E.M.; *n* = 7 (Tie2-control + isotype control), 9 (Tie2-control + anti-TNF Ab), 7 (Tie2-L613V + isotype control) or 5 (Tie2-L613V + anti-TNF Ab); with 200 CMs counted per sample; \**P* < 0.05, Bonferroni’s post-test when ANOVA was significant; # *P* < 0.05, two-tailed Student’s *t* test). **d**, Ejection fraction and **e**, fractional shortening, measured by echocardiography, showing preserved cardiac function following TNF inhibition. **f**, Models illustrating how the combined cell-autonomous and non cell-autonomous actions of an activating RAF1 mutants cause NS-cardiomyopathy (see text for details).

## Discussion

We find that combinatorial interactions of CMs, ECs and cardiac fibroblasts underlie the pathogenesis of RAF1 mutant NS-associated cardiomyopathy (Fig. 6f). In line with the conventional view, altered cardiac contractility in this setting arises from a CM-intrinsic defect(s) that causes increased sensitivity of the cardiac contractile apparatus to Ca^2+^. But although CM-specific mutant RAF1 expression slightly increases ventricular wall thickness, the major pro-hypertrophic signals, including TNF (and IL6), emanate from mutant ECs. Moreover, aberrant RAF1 activity in CMs or cardiac fibroblasts, but not ECs, contributes to pressure overload-induced fibrosis in NS cardiomyopathy.

Our understanding of the role of RAS/ERK pathway in the heart has been based largely on transgenic over-expression or knockout mouse models, typically with a restricted focus on CMs^36–41^. For example, transient overexpression of a NS-associated *PTPN11* gain-of-function mutation in rat CMs increases Ca^2+^ oscillatory frequency^42^. CMs isolated from a transgenic mouse model of LVH associated with another RASopathy, NS with multiple lentigines (NS-ML), exhibit increased calcium transients and SERCA expression^43^. CMs derived from *BRAF* mutant human induced pluripotent cells (hiPSCs) from patients with cardio-facial-cutaneous syndrome also have increased calcium transients^44^. Conversely, we find that a NS-associated activating RAF1 mutant, when expressed solely in CMs, does not affect Ca^2+^ levels, but instead, increases the Ca^2+^ sensitivity of the contractile apparatus. We observe a similar alteration in calcium sensitivity in RAF1 mutant human CMs derived from hiPSCs or ESCs (T. Araki, unpublished observations). Precisely how mutant RAF1 regulates Ca^2+^ sensitivity remains to be elucidated, although it likely involves altered phosphorylation of key calcium handling or myofilament proteins^45,46^. Taken together, though, these finding suggest that different RASopathy mutations might alter CM contractility in distinct ways.

Surprisingly, mutant RAF1 expression in CM causes minimal hypertrophy. By contrast, EC-restricted mutant expression does not affect contractility, but evokes hypertrophy. Excess TNF produced by mutant ECs can, via a cytokine hierarchy that includes IL6 and probably other agonists, account for their pro-hypertrophic effects *in vitro* (i.e., co-culture/Transwell assays). Interfering with TNF action (by neutralizing antibody injections) also normalizes the increased ventricular wall thickness and CM size *in vivo*. Reverse remodeling is not complete, however, as chamber size and HW/BW remain elevated. Notably, complete remodeling is possible, as MEK inhibitor treatment fully reverses pathological hypertrophy in global *Raf1^L613V/+^* mice^22^.

Failure of anti-TNF therapy to restore normal cardiac chamber size presumably reflects other, TNF-independent, paracrine signals from mutant ECs. Matrix metalloproteinases (MMPs), tissue inhibitor of metalloproteinases (TIMPs) and ADAMs are implicated in pathological cardiac hypertrophy^47,48^, and several of these are dysregulated in RAF1 mutant ECs (Supplementary Fig. 7d). Although TNF stimulates production of some MMPs, most MMP promoters contain AP-1 and/or PEA3 elements and could be stimulated directly by ERK-catalyzed phosphorylation^47^. Alternatively, TNF could have a “Goldilocks” effect, with too much or too little having deleterious consequences, and our anti-TNF therapy might have been insufficient to restore normal cardiac size. Indeed, increased TNF levels are seen in patients with HCM and other cardiac disorders^49–55^. However, results from large clinical trials, as well as several case reports, suggest that too much TNF inhibition can lead to dilated cardiomyopathy and/or heart failure^56–59^. Notably, cardiac function is preserved in our TNF antibody-treated mice. Nevertheless, future studies will be needed to determine the effect of TNF inhibition on all of the paracrine signaling molecules altered in RAF1 mutant ECs and to assess the dose-response relationship between TNF inhibition and cardiac hypertrophy.

Increasing evidence points to the importance of communication between myocytes and non-myocytes in modulating cardiac function and structure under physiological and pathological conditions^31,60,61^. Although ECs clearly play essential roles in cardiac valve development and disease, including defects associated with RAS/ERK pathway dysregulation^62,63^, the role of ECs in driving cardiac hypertrophy has been less explored. While our manuscript was in revision, Lauriol *et al*. reported that NS-ML-associated hypertrophy arises from expression of catalytically impaired SHP2 (*PTPN11*) solely in EC^64^. They observed ventricular thinning and delayed septal closure in developing hearts, which they attributed to increased AKT and decreased FOXP1 and NOTCH1 signaling. Adult NS-ML mice, however, have cardiac hypertrophy^65^, which Lauriol *et al*. suggest might reflect a compensatory response to ventricular thinning during development. Importantly, Tie2-L613V embryos do not exhibit ventricular thinning. Furthermore, it seems unlikely that a developmental defect fully accounts for the cardiac hypertrophy in NS-ML mice, as postnatal rapamycin treatment rescues NS-ML-associated hypertrophy, suggestive of ongoing pro-hypertrophic signaling^65^. Very recently, Josowitz *et al*. reported that fibroblast-like cells derived from *BRAF* mutant hiPSCs have pro-hypertrophic effects on hiPSC-derived CMs, via paracrine transforming growth factor β (TGFβ) production^44^. Whether mutant CFs promote hypertrophy in the whole organism context remains to be determined, although conceivably, distinct cardiac cell types are responsible for pathological hypertrophy in different RASopathies.

Regardless, our results, along with other recent work^44,60,64^, argue for a paradigm shift away from the myocyte-centric view of cardiac development and disease. Aberrant EC signaling is likely a recurring pathogenic mechanism in RASopathy-associated cardiomyopathy, and might well contribute to other genetic or even secondary forms of LVH. Consequently, it will be important to test whether ECs is/are a/the source of TNF and IL6 in the more common types of pathological LVH. Furthermore, anti-TNF and/or IL6 signaling pathway agents might prove beneficial as targeted therapy for treating RASopathy patients with severe LVH.

## Methods

### Generation of mice

Mlc2v-L613V, Fsp1-L613V, Tie2-L613V, Nfatc1-L613V, cTnT-L613V mice were generated by crossing inducible *Raf1^L613Vfl/+^* knock-in mice (129Sv × C57BL/B6)^22^ and Mlc2v-Cre^23^, Tie2-Cre^66^, Nfatc1-Cre^67^, Fsp1-Cre^68^ or cTnT-Cre^24^, respectively. Sixteen-week old male mice were used for all cardiac physiology experiments. For all experiments, investigators were blinded to animal genotype. PCR genotyping was performed as described^22^; detailed conditions are available from J.C.Y. All animal studies were approved by the Animal Care Committees of University Health Network and the University of Guelph and were performed in accordance with the standards of the Canadian Council on Animal Care. Based on previous results^22^, we had 80% power to detect differences of >15% in all cardiac parameters with at least 8 mice per group with a significance level of 0.05 (Student’s *t* test).

### Echocardiographic and hemodynamic Analysis

For echocardiographic and hemodynamic analysis, mice were anesthetized with isoflurane/oxygen (2%/100%). Echocardiography was performed with the Vevo2100 system (VisualSonics Inc., Toronto, ON, CA) and the MS550D transducer at 40 MHz. Acquired M-mode images were analyzed with the LV-trace function from the cardiac package (VisualSonics Inc., Toronto, ON, CA). All measurements were made between 12 and 5pm. Following echocardiographic assessment, mice were transferred to a warmed surgical plate, maintained at 37°C using a heating lamp and rectal probe, and a 1.2F catheter (FTS-1211B-0018; Scisense Inc.) was inserted via the right carotid into the LV. Hemodynamic signals were digitized at 2000 Hz and recorded using iWorx^®^ analytic software (Labscribe2, Dover, NH, USA). Following data collection, mice were sacrificed by cardiac excision.

### Primary cell isolation and culture

Adult ventricular CMs were isolated from male Mlc2v-control or -L613V mice between 2-5 months of age. After heparin was administered i.p. (10 IU/g body weight), mice were deeply anesthetized with isofluorane, and their hearts were removed and placed in ice-cold modified Ca^2+^ free Tyrode’s solution (pH 7.3). Excised hearts were perfused with the same solution at 37°C for 8 min, after which Type II collagenase (88 U/ml, Worthington Biochemical Corporation) plus pure Yakult Collagenase (125 U/ml, Yakult Pharmaceutical Industry Corporation) were introduced for 10-12 min. The inner layer of the left ventricle was removed into a Ca^2+^-free Tyrode’s solution, gently cut into small pieces, and triturated using a polished glass pipette. Dissociated CMs were allowed to settle for 5 min in a 50ml conical tube, the supernatant was discarded to remove residual enzyme, and the CMs were re-suspended. This settling/resuspension sequence was repeated twice. Isolated CMs were stored at room temperature (25 °C) until use.

To isolate neonatal CMs, single cell suspensions were prepared using the Neonatal Heart Dissociation Kit (Miltenyi Biotec), according to the manufacturer’s instructions. Non-myocytes were depleted by magnetic activated cell sorting (MACS; Miltenyi Biotec).

For non-myocyte isolation, postnatal day 4-7 hearts were minced into small pieces. Single cell suspensions were prepared by enzymatic digestion with collagenase (300 U/ml) and hyaluronidase (100 U/ml) from STEMCELL Technologies Inc. for 45 minutes with gentle rotation, followed by mechanical dissociation. Endothelial/endocardial cells and fibroblasts were enriched by MACS using CD31 and CD90 microbeads, respectively.

Isolated neonatal CMs or postnatal non-myocytes were plated onto 2% gelatin-coated 96-well plates (day 0), and maintained or co-cultured in StemPro-34 media (Life Technologies) supplemented with rhbFGF (5 ng/ml; R&D) and rhVEGF (20 ng/ml; Cedarlane). For dose-response assays, wild type CMs were treated with mouse recombinant TNF or IL6 (ThermoFisher) for three days at the indicated doses. Co-cultured cells were washed with HBSS once before a media change on day 1, at which point cells were subjected to neutralizing antibody treatments, as indicated. For Transwell assays, CMs were plated on the gelatin-coated bottoms of the Transwell system (Corning Costar), and CD31^+^ cells were plated onto collagen-coated membrane inserts.

### Measurement of Ca^2+^ transients

Aliquots of freshly isolated ventricular CMs were placed in a storage solution containing 1 μM Fura-2-acetoxymethyl ester (AM) for 20 min at room temperature (25 °C), after which the cells were dispersed onto a glass-bottomed chamber equipped with platinum electrodes to allow field stimulation. After 3 min, myocytes were washed for 5 min with a modified Krebs-Henseleit perfusion solution (120 mM NaCl, 5.4 mM KCl, 1 mM MgCl2, 10 mM HEPES, 10 mM Glucose, 19 mM NaHCO3, 1.2 mM CaCl2 and 1 mM Na pyruvate; pH 7.3). Fura-2 fluorescence was measured by illuminating CMs via the rear light port of an Olympus IX70 microscope, alternating every 10 seconds between 340 nm or 380 nm light (10 nm band-pass) originating from a 75 W xenon arc lamp. The light was projected to the CMs by a 40X objective (UApo/340, 40X /0.9NA, Olympus). The emitted light at 510 nm (±20 nm) from the CMs was projected onto an Evolve 128 camera (Photometrics) and acquired at 100 frames/s via Metamorph software (Molecular Devices). Images were analyzed using ImageJ. Field stimulation was applied for 5 min at 0.5 Hz, using 2 ms pulses at twice the threshold voltage (typically 10 V/cm). Only rod-shaped CMs displaying clear striation patterns and stable shortening patterns were used for Ca^2+^ transient recordings. To determine basal auto-fluorescence, a Kreb’s-Henseleit solution containing 2 mM Mn^2+^ (and no Ca^2+^) was applied, which quenches the Fura-2 fluorescence signal. Ca^2+^ transients were quantified by the equation:

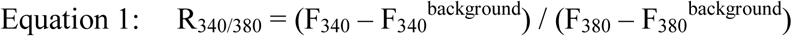
where F_340_ and F_380_ are the intensities of the 510 nm fluorescence originating from the Fura-2 in the CMs when illuminated with light at 340 nm and 380 nm, respectively. F_340_^background^ and F_380_^background^ are the fluorescence levels measured in the same region of the CMs after Mn^2+^ addition.

### Measurement of force and force-Ca^2+^ relationships

Freshly isolated ventricular CMs were loaded with Fura-2 as described above. CMs were attached to a pair of glass rods using the biocompatible adhesive MyoTak (World Precision Instruments, Inc.), as per the manufacturer’s directions. After “adhesion”, CMs were lifted from the glass and were field stimulated at various rates (0.1-10 Hz) using 2 ms pulses at twice the threshold voltage (typically 10 V/cm). During recording of force and Ca^2+^ transients myocytes were illuminated using red light (650-690 nm). Light (510 nm plus 650-690 nm) was collected with a 40X objective (UApo/340, 40X /0.9NA, Olympus) on a IX 70 Olympus microscope and projected through a dichroic mirror, which reflected the 510 nm light to the Photometrics camera and passed the red light to the a MyocamS Camera (IonOptix). Force is measured as pixels of deflection of the glass rods arising from CM contraction in response to field stimulation. The force was normalized to CM cross-sectional area (CSA), which was determined at the end of the experiment by imaging each cell with its long axis aligned with the optical axis of the microscope. This approach allows the width and thickness of CMs to be measured accurately. CSA was estimated using the equation π*1/2width*1/2thickness, as suggested^69^. The images of the cells generated by the red light were also analyzed by fast Fourier-transforming selected regions of the CMs. This algorithm allows sarcomere length to be rapidly estimated during contractions. Force (in pixels) and Ca^2+^ (R_340/380_) were recorded while keeping the diastolic sarcomere length of the CMs at 1.8 μm.

### Transverse aortic constriction (TAC)

Mice (8-9 week old, 24-32 g) were anesthetized with isoflurane/oxygen (2%: 100%), intubated, and ventilated (Harvard Apparatus) at 150-180 breaths/minute, 250 μL/breath. The transverse aorta was constricted with a 7/0 silk suture tied around a 27-gauge needle, as described^22^. Pressure overload was maintained for 2 wks. Mice were housed on a 12- hour light/dark cycle (8am/8pm) and monitored twice daily.

### Cytokine measurements

Conditioned media were collected from cultured cells two days after a media change and immediately stored at -80 °C until analysis. Luminex bead-based multiplex cytokine arrays (Millipore) were performed in magnetic plates, according to the manufacturer’s instructions, and read with a Luminex 100 Reader. Data were analyzed using Bio-plex Manager 6.0. IL6 levels following anti-TNF antibody treatment were assessed using a commercially available ELISA (Sigma), according to the manufacturer’s instructions. All assays were performed in duplicate.

### qRT-PCR

Tissues were flash frozen in liquid nitrogen, and cells were washed with ice-cold HBSS before extraction with QIAzol lysis reagent (Qiagen). RNA was isolated using miRNeasy (Qiagen), according to the manufacturer’s instructions. cDNA was synthesized by reverse transcription using oligo(dT) primers and the Superscript III First Strand Synthesis System (Invitrogen). Transcripts were detected and quantified by qRT-PCR using the QuantiFast SYBR Green PCR Kit (Qiagen). Primer sequences are provided in Supplementary Table 3. All values were normalized to *Gapdh* levels.

### RNA sequencing

RNA (150 ng) from each sample was reversed transcribed using the Illumina TruSeq Stranded mRNA kit. cDNA libraries were sized on an Agilent Bioanalyzer, and their concentrations were validated by qPCR. Four different libraries were normalized to 10nM and pooled, 8 pM of pooled libraries were loaded onto an Illumina cBot for cluster generation, and the flow cell was subjected to 100-cycles of paired-end sequencing on an Illumina HighSeq 2000. normalized and pooled together, and loaded onto Illumina cBot for cluster generation. Pair-end sequencing (100 cycles) was performed on an Illumina HiSeq 2000. The raw sequence data, in the form of FASTQ files, was aligned to the mouse genome (UCSC mm10, iGenome GTF definition file) using the BOWTIE/TOPHAT pipeline (BOWTIE v2.2.3, TOPHAT 2.0.13). Accessory programs for the alignment stage include cutadapt (1.7.1). Transcripts were assembled and gene expression levels were estimated with Cufflinks (v.2.2.1) using iGenome GTF file. Cufflinks output result was load into R (3.1.1)/cummeRbund (2.6.1) for final output of results and graphing.

### Histology

Hearts were harvested in the relaxed state by perfusion-fixation with 1% KCl in PBS, followed by 10% buffered formalin overnight. Staining with H&E, Picro Sirius Red (PSR; 500ml of saturated picric acid solution and 0.5 g of Direct Red 80 from Sigma Aldrich) and Masson-trichrome was performed on paraffin-embedded sections (5 μm), according to standard practice. FITC-conjugated wheat germ agglutinin (WGA; Sigma-Aldrich) was used with appropriate antibodies and DAPI for the approximation of cross-sectional area in CMs with centrally located nuclei (∼200/sample). ImageJ software was used for quantification. For BrdU incorporation assays, pregnant mice at E16.5 were subjected to BrdU i.p. injection (100 μg/g body weight) one hour before sacrifice. Embryos were fixed overnight in 10% buffered formalin and embedded in paraffin. Sections were stained with primary antibodies against BrdU (1:50 Abcam), followed by secondary antibodies conjugated with F(ab)2 biotin (1:500; Research Diagnostics Inc.), and developed using the Vectastain Elite ABC Kit (Vector Laboratories). All sections were counterstained with hematoxylin. BrdU-positive cells were counted in 20 randomly selected fields from each sample.

For immunofluorescence staining of primary cultures, cells were washed with 1% KCl in ice-cold PBS three times before fixation with 4% paraformaldehyde for 15 minutes. Cells were stained with 5 μg/ml monoclonal rat anti-mouse CD31 antibody (Clone MEC13.3; BD Pharmingen) or 5 μg/ml monoclonal rat anti-mouse CD90 antibody (Clone 30-H12; Biolegend) and, after permeabilization with 1% Triton X-100, 2 μg/ml monoclonal mouse anti-mouse cTnT antibody (Clone 13-11; Thermo Scientific) and DAPI. Primary antibody binding was visualized by using fluorescence-coupled anti-rat or anti-mouse secondary antibodies at 1:250 dilutions. Images were taken by scanning across the well, with all visualized CMs (at least 300) counted per well.

### Immunoblotting

Hearts from 10 to 16-week old mice were harvested and flash frozen in liquid nitrogen. For Ca^2+^ handling proteins, total protein extracts from hearts were prepared by homogenization in 1% SDS (50 mM Tris-HCl, pH 8, 100 mM NaCl, 2 mM EDTA). For signaling proteins, total protein extracts from hearts were prepared by homogenization in RIPA buffer (50 mM Tris-HCl, pH 7.5, 150 mM NaCl, 2 mM EDTA, 1% NP-40, 0.5% Na deoxycholate, and 0.1% SDS), containing a protease and phosphatase inhibitor cocktail (40 μg/ml PMSF, 20 mM NaF, 1 mM Na_3_VO_4_, 10 mM β-glycerophosphate, 10 mM sodium pyrophosphate, 2 μg/ml antipain, 2 μg/ml pepstatin A, 20 μg/ml leupeptin, and 20 μg/ml aprotinin). Lysates were centrifuged at 16,100 g for 15 minutes at 4 °C. Clarified supernatants (70 μg for detection of pSTAT3 and pSTAT5; 10-25 μg for others) were resolved by SDS-PAGE and analyzed by immunoblotting. Antibodies used for immunoblots included: anti-ERK2 (7 ng/ml, Clone D-2; Santa Cruz Biotechnology Inc.), and anti-SERCA2 (Clone D51B11; #9580), anti-phospho- p44/42 MAPK (#9101), -phospho-MEK1/2 (#9121), -phospho-AKT (S473, #9271), -phospho-AKT (T308, Clone 244F9, #4056), -AKT1 (Clone C73H10, #2938), -STAT3 (#9132), -phospho- STAT3 (Y705, #9131), -STAT5 (#9163), -phospho-STAT5 (Y694, #9351; all at 1:1000 dilutions from Cell Signaling Technology). Primary antibodies were visualized by IRDye infrared secondary antibodies (1:15,000 dilution for 680LT anti-mouse IgG and 1:10,000 dilution for 800CW anti-rabbit IgG), using the Odyssey Infrared Imaging System (LI-COR Biosciences). Immunoblot signals were quantified by using Odyssey version 3.0 software.

### Flow Cytometry

CD31^+^ cells were stained with 2 μg/ml PE-conjugated anti-mouse CD31 (Clone MEC13.3; BD Pharmingen) and 2 μg/ml APC-conjugated anti mouse CD45 (Clone 30F11; BioLegend) and analyzed on a LSR II flow cytometer (BD Bioscience). Flow cytometric data were analyzed with FlowJo software (TreeStar).

### Anti-TNF neutralizing antibody experiments

LEAF^™^ purified anti-mouse TNF antibody (MP6-XT22, BioLegend) or LEAF^™^ purified rat IgG1 isotype control (RTK2071, BioLegend) was injected i.p. (5 mg/kg body weight) twice weekly for the indicated times.

### Statistical analysis

All experiments were performed on biological replicates unless otherwise specified. The number of biological replicates is represented by “n”. Sample size for each experimental group/condition is reported in the appropriate figure legends and methods. For cell culture experiments, sample size was not pre-determined, and all samples were included in analysis. All data are presented as mean ± standard error of the means (S.E.M.). For normally distributed data, differences between two groups were evaluated by Student’s *t* test and differences between multiple groups were evaluated by ANOVA, followed by Bonferroni’s post-test. For non-normally distributed data (Fig. 5b, c, f, and g), differences between two groups were evaluated with the non-parametric Mann-Whitney test and differences between multiple groups were evaluated by the non-parametric Kruskal–Wallis one-way analysis of variance (ANOVA), followed by Dunn’s post-hoc test. For all experiments, except for cell culture experiments (Fig. 5b, c, f, and g), the between-group variances were similar and data were symmetrically distributed. All analyses and graphs were generated with GraphPad Prism 5. A p-value of <0.05 was considered significant.

## Data availability

The raw RNAseq data have 575 been deposited in GEO under accession code GSE95739. All other data supporting the findings of this study are available from the corresponding authors on request.

## End notes

### Acknowledgements

We thank Drs. G. Keller, M. Sherrid, G. Fishman and M. Feldmann for helpful comments, L. Morikawa and N. Law for assistance with histology, J. Tsao, Z. Lu and C. Virtanen for help with RNAseq, A. Sayad for other statistical analyses, and B. Gurbaksh and P. Yao for help with Luminex bead-based cytokine array analysis. This work was supported by R01 HL 083273 (B.G.N.) and Canadian Institutes of Health Research grants MOP111159 (J.A.S.) and 106526 to (T.A.) and MOP-83453 to (P.H.B.). B.G.N. was a Canada Research Chair, Tier 1, and work in his laboratory was supported by grants from the Ontario Ministry of Health and Long Term Care and The Princess Margaret Cancer Foundation. J.A.S. is a new investigator with the Canadian Heart and Stroke Foundation, and J.C.Y. was supported by a CIHR CGS-D scholarship.

### Author Contributions

B.G.N., X.W., J.A.S and T.A. conceived the project. B.G.N., T.A., P.H.B. and J.A.S. supervised the research. J.C.Y., P.H.B., T.A., J.A.S., and B.G.N. designed the experiments. J.C.Y., M.J.P., X.T., X.W. and J.A.S. performed the experiments. All authors participated in data analysis. J.C.Y. and B.G.N. wrote the manuscript with the help of all of the authors.

### Competing Financial Interest

The authors declare no competing financial interests.

### Author Information

Reprints and permissions information are available at www.nature.com/reprints. Readers are welcome to comment on the online version of the paper. Correspondence and requests for materials should be addressed to B.G.N., T.A., or J.A.S.

## References

1. Maron, B. J. & Maron, M. S. Hypertrophic cardiomyopathy. Lancet 381, 242–255 (2013).

2. Maron, B. J. Hypertrophic Cardiomyopathy: A Systemic Review. JAMA 287, 1308–1320 (2002).

3. Frey, N., Luedde, M. & Katus, H. a. Mechanisms of disease: hypertrophic cardiomyopathy. Nat. Rev. Cardiol. 9, 91–100 (2011).

4. Drazner, M. H. The progression of hypertensive heart disease. Circulation 123, 327–334 (2011).

5. Seidman, J. G. & Seidman, C. The Genetic Basis for Cardiomyopathy. Cell 104, 557–567 (2001).

6. Rader, F., Sachdev, E., Arsanjani, R. & Siegel, R. J. Left Ventricular Hypertrophy in Valvular Aortic Stenosis: Mechanisms and Clinical Implications. Am. J. Med. 128, 344–352 (2015).

7. Alcalai, R., Seidman, J. G. & Seidman, C. E. Genetic basis of hypertrophic cardiomyopathy: From bench to the clinics. J. Cardiovasc. Electrophysiol. 19, 104–110 (2008).

8. Rohini, A., Agrawal, N., Koyani, C. N. & Singh, R. Molecular targets and regulators of cardiac hypertrophy. Pharmacol. Res. 61, 269–280 (2010).

9. Heineke, J. & Molkentin, J. D. Regulation of cardiac hypertrophy by intracellular signalling pathways. Nat. Rev. Mol. Cell Biol. 7, 589–600 (2006).

10. Dorn 2nd, G. W., Force, T. & Ii, G. W. D. Protein kinase cascades in the regulation of cardiac hypertrophy. J Clin Invest 115, 527–537 (2005).

11. Sala, V. & Gallo, S. Signaling to Cardiac Hypertrophy: Insights from Human and Mouse RASopathies. Mol. Med. 18, 1 (2012).

12. Tidyman, W. E. & Rauen, K. a. The RASopathies: developmental syndromes of Ras/MAPK pathway dysregulation. Curr. Opin. Genet. Dev. 19, 230–236 (2009).

13. Rauen, K. a. The RASopathies. Annu. Rev. Genomics Hum. Genet. 14, 355–69 (2013).

14. Gelb, B. D. & Tartaglia, M. RAS signaling pathway mutations and hypertrophic cardiomyopathy: Getting into and out of the thick of it. J. Clin. Invest. 121, 844–847 (2011).

15. Cirstea, I. C. et al. A restricted spectrum of NRAS mutations causes Noonan syndrome. Nat. Genet. 42, 27–29 (2009).

16. Flex, E. et al. Activating mutations in RRAS underlie a phenotype within the RASopathy spectrum and contribute to leukaemogenesis. 23, 4315–4327 (2014).

17. Cordeddu, V. et al. Mutation of SHOC2 promotes aberrant protein N-myristoylation and causes Noonan-like syndrome with loose anagen hair. Nat. Genet. 41, 1022–1026 (2009).

18. Yamamoto, G. L. et al. Rare variants in SOS2 and LZTR1 are associated with Noonan syndrome. J. Med. Genet. 1–9 (2015). doi:10.1136/jmedgenet-2015-103018

19. Chen, P.-C. et al. Next-generation sequencing identifies rare variants associated with Noonan syndrome. Proc. Natl. Acad. Sci. U. S. A. 111, 11473–11478 (2014).

20. Gripp, K. W. et al. A Novel Rasopathy Caused by Recurrent De Novo Missense Mutations in PPP1CB Closely Resembles Noonan Syndrome with Loose Anagen Hair. Am J Med Genet 170A, 2237–2247 (2016).

21. Pandit, B. et al. Gain-of-function RAF1 mutations cause Noonan and LEOPARD syndromes with hypertrophic cardiomyopathy. Nat. Genet. 39, 1007–1012 (2007).

22. Wu, X. et al. MEK-ERK pathway modulation ameliorates disease phenotypes in a mouse model of Noonan syndrome associated with the Raf1L613V mutation. J. Clin. Invest. 121, 1009–1025 (2011).

23. Chen, J., Kubalak, S. W. & Chien, K. R. Ventricular muscle-restricted targeting of the RXRalpha gene reveals a non-cell-autonomous requirement in cardiac chamber morphogenesis. Development 125, 1943–1949 (1998).

24. Jiao, K. et al. An essential role of Bmp4 in the atrioventricular septation of the mouse heart service An essential role of Bmp4 in the atrioventricular septation of the mouse heart. Genes Dev. 17, 2362–2367 (2003).

25. Backx, P. H., Gao, W. D., Azan-Backx, M. D. & Marban, E. The relationship between contractile force and intracellular [Ca2+] in intact rat cardiac trabeculae. J. Gen. Physiol. 105, 1–19 (1995).

26. Kentish, J. C. & Wrzosek, A. Changes in force and cytosolic Ca ¥ concentration after length changes in isolated rat ventricular trabeculae. J. Physiol. 506, 431–444 (1998).

27. Iwano, M. et al. Evidence that fibroblasts derive from epithelium during tissue fibrosis. J. Clin. Invest. 110, 341–350 (2002).

28. Moore-morris, T. et al. Resident fi broblast lineages mediate pressure overload – induced cardiac fi brosis. 124, 1–14 (2014).

29. Kong, P., Christia, P., Saxena, A., Su, Y. & Frangogiannis, N. G. Lack of specificity of fibroblast-specific protein 1 in cardiac remodeling and fibrosis. Am. J. Physiol. Heart Circ. Physiol. 305, H1363–72 (2013).

30. Zeisberg, E. M. et al. Endothelial-to-mesenchymal transition contributes to cardiac fibrosis. Nat. Med. 13, 952–961 (2007).

31. Brutsaert, D. L. Cardiac endothelial-myocardial signaling: its role in cardiac growth, contractile performance, and rhythmicity. Physiol. Rev. 83, 59–115 (2003).

32. Newell, C. L., Deisseroth, a B. & Lopezberestein, G. Interaction of Nuclear Proteins With an Ap-1 Cre-Like Promoter Sequence in the Human Tnf-Alpha Gene. J. Leukoc. Biol. 56, 27–35 (1994).

33. Kramer, B., Wiegmann, K. & Kronke, M. Regulation of the human TNF promoter by the transcription factor Ets. Journal of Biological Chemistry 270, 6577–6583 (1995).

34. Feldmann, M.& Maini, R. N. Anti-TNF (alpha) therapy or rheumatoid arthritis□: What have we learned□? Annu. Rev. Immunol. 19, 163–96 (2001).

35. Neurath, M. F. Cytokines in inflammatory bowel disease. Nat. Rev. Immunol. 14, 329 – 342 (2014).

36. Hunter, J. J. et al. Nucleic Acids, Protein Synthesis, and Molecular Genetics□: Ventricular Expression of a MLC-2v- ras Fusion Gene Induces Cardiac Hypertrophy and Selective Diastolic Dysfunction in Transgenic Mice Ventricular Expression of a MLC-2v- ras Fusion Gene Induces. J. Biol. Chem. 270, 23173–23178 (1995).

37. Purcell, N. H. et al. Genetic inhibition of cardiac ERK1/2 promotes stress-induced apoptosis and heart failure but has no effect on hypertrophy in vivo. Proc. Natl. Acad. Sci. 104, 14074–14079 (2007).

38. Harris, I. S. et al. Raf-1 kinase is required for cardiac hypertrophy and cardiomyocyte survival in response to pressure overload. Circulation 110, 718–723 (2004).

39. Zheng, M. et al. Sarcoplasmic reticulum calcium defect in Ras-induced hypertrophic cardiomyopathy heart. Am J Physiol Hear. Circ Physiol 286, H424–33 (2004).

40. Bueno, O. F. et al. The MEK1-ERK1/2 signaling pathway promotes compensated cardiac hypertrophy in transgenic mice. EMBO J. 19, 6341–50 (2000).

41. Yamaguchi, O. et al. Cardiac-specific disruption of the. October 114, (2004).

42. Uhlen, P. et al. Gain-of-function/Noonan syndrome SHP-2/Ptpn11 mutants enhance calcium oscillations and impair NFAT signaling. Proc. Natl. Acad. Sci. 103, 2160–2165 (2006).

43. Clay, S. A., Domeier, T. L., Hanft, L. M., Mcdonald, K. S. & Krenz, M. Elevated Ca 2 □ transients and increased myofibrillar power generation cause cardiac hypercontractility in a model of Noonan syndrome with multiple lentigines. 1086–1095 (2015). doi:10.1152/ajpheart.00501.2014

44. Josowitz, R. et al. Autonomous and non-autonomous defects underlie hypertrophic cardiomyopathy in BRAF-mutant hiPSC-derived cardiomyocytes. Stem Cell Reports 7, 355–369 (2016).

45. Layland, J., Solaro, R. J. & Shah, A. M. Regulation of cardiac contractile function by troponin I phosphorylation. 66, 12–21 (2005).

46. Yuan, C. & Solaro, R. J. Myofilament proteins: From cardiac disorders to proteomic changes. Proteomics - Clin. Appl. 2, 788–799 (2008).

47. Spinale, F. G. Myocardial Matrix Remodeling and the Matrix Metalloproteinases□: Influence on Cardiac Form and Function. 1285–1342 (2007). doi:10.1152/physrev.00012.2007.

48. Olivetti, G., Capasso, J. M., Sonnenblick, E. H. & Anversa, P. Side-to-Side Slippage of Myocytes Participates in Ventricular Wall Remodeling Acutely After Myocardial Infarction in Rats.

49. Högye, M., Mándi, Y., Csanády, M., Sepp, R. & Buzás, K. Comparison of circulating levels of interleukin-6 and tumor necrosis factor-alpha in hypertrophic cardiomyopathy and in idiopathic dilated cardiomyopathy. Am. J. Cardiol. 94, 249–251 (2004).

50. Patel, R. et al. Variants of trophic factors and expression of cardiac hypertrophy in patients with hypertrophic cardiomyopathy. J. Mol. Cell. Cardiol. 32, 2369–2377 (2000).

51. Velten, M. et al. Priming with synthetic oligonucleotides attenuates pressure overload-induced inflammation and cardiac hypertrophy in mice. Cardiovasc. Res. 96, 422–432 (2012).

52. Bautista, L. E., Vera, L. M., Arenas, I. a & Gamarra, G. Independent association between inflammatory markers (C-reactive protein, interleukin-6, and TNF-alpha) and essential hypertension. J. Hum. Hypertens. 19, 149–154 (2005).

53. Vázquez-Oliva, G., Fernández-Real, J. M., Zamora, a, Vilaseca, M. & Badimón, L. Lowering of blood pressure leads to decreased circulating interleukin-6 in hypertensive subjects. J. Hum. Hypertens. 19, 457–462 (2005).

54. Lee, D. L. et al. Angiotensin II hypertension is attenuated in interleukin-6 knockout mice. Am. J. Physiol. Heart Circ. Physiol. 290, H935–H940 (2006).

55. Levine, B., Kalman, J., Mayer, Lloyd, Fillit, H.M., Packer, M. Elevated circulating levels of tumor necrosis factor in severe chronic heart failure. N. Engl. J. Med. (1990).

56. Mann, D. L. et al. Targeted Anticytokine Therapy in Patients With Chronic Heart Failure: Results of the randomized ethanercept worldwide evaluation (RENEWAL). Circulation 109, 1594–1603 (2004).

57. Rocha, F. A. C. & Silva, F. S. Reversible Heart Failure in a Patient Receiving Etanercept for Aankylosing Spondylitis. J Clin Rheumatol 16, 81–82 (2010).

58. Ma, K., Dormand, H., Neyses, L. & Ma, M. Heart Failure with Etanercept Therapy □: A Case Report. J Clin Exp Cardiol. 4, 3–5 (2013).

59. Emmert, M. Y. et al. Severe cardiomyopathy following treatment with the tumour necrosis factor- a inhibitor adalimumab for Crohn’ s disease. Eur. J. Heart Fail. 11, 1106–1109 (2009).

60. Tian, Y. & Morrisey, E. E. Importance of myocyte-nonmyocyte interactions in cardiac development and disease. Circ. Res. 110, 1023–1034 (2012).

61. Takeda, N. & Manabe, I. Cellular Interplay between Cardiomyocytes and Nonmyocytes in Cardiac Remodeling. Int. J. Inflam. 2011, 535241 (2011).

62. Gitler, A. D. et al. Nf1 has an essential role in endothelial cells. Nat. Genet. 33, 75–79 (2003).

63. Araki, T. et al. Noonan syndrome cardiac defects are caused by PTPN11 acting in endocardium to enhance endocardial-mesenchymal transformation. Proc. Natl. Acad. Sci. U. S. A. 106, 4736–4741 (2009).

64. Lauriol, J. et al. Developmental SHP2 dysfunction underlies cardiac hypertrophy in Noonan syndrome with multiple lentigines. J Clin Invest 8126, 2989–3005 (2016).

65. Marin, T. M. et al. Rapamycin reverses hypertrophic cardiomyopathy in a mouse model of LEOPARD syndrome – associated PTPN11 mutation. 121, (2011).

66. Koni, P. a et al. Conditional vascular cell adhesion molecule 1 deletion in mice: impaired lymphocyte migration to bone marrow. J. Exp. Med. 193, 741–754 (2001).

67. Wu, B. et al. Endocardial cells form the coronary arteries by angiogenesis through myocardial-endocardial VEGF signaling. Cell 151, 1083–1096 (2012).

68. Bhowmick, N. a et al. TGF-beta signaling in fibroblasts modulates the oncogenic potential of adjacent epithelia. Science 303, 848–851 (2004).

69. King, N. M. P. et al. Mouse intact cardiac myocyte mechanics□: cross-bridge and titin-based stress in unactivated cells The Journal of General Physiology. J. Gen. Physiol. 137, 81–91 (2010).

